# The glucocorticoid dexamethasone impairs the expression of anti-viral mediators in activated macrophages by inhibiting both expression and function of interferon β

**DOI:** 10.1101/2023.05.03.539212

**Authors:** John D O’Neil, Oliwia O Bolimowska, Sally A Clayton, Tina Tang, Kalbinder K Daley, Samuel Lara-Reyna, Jordan Warner, Claire S Martin, Rahul Y Mahida, Rowan S Hardy, J Simon C Arthur, Andrew R Clark

## Abstract

Glucocorticoids potently inhibit expression of many inflammatory mediators, and have been very widely used to treat both acute and chronic inflammatory diseases for more than seventy years. However, they can have several unwanted effects, amongst which immunosuppression is one of the most common. Here we investigated effects of the synthetic glucocorticoid dexamethasone on the responses of primary mouse bone marrow-derived macrophages to the pro-inflammatory agonist lipopolysaccharide (LPS). At the mRNA level, dexamethasone inhibited the LPS-induced expression of more than 100 genes that are involved in cell-intrinsic defence against viral pathogens. Expression of most of the corresponding proteins was also reduced by dexamethasone. This antiviral disarmament occurred at two distinct levels. First, dexamethasone strongly and dose-dependently inhibited the expression of the type I interferon IFNβ by LPS-activated macrophages. IFNβ mediates an autocrine positive feedback loop in LPS-treated macrophages, promoting the expression of antiviral genes and other interferon-stimulated genes. Hence reduction of IFNβ expression contributes to impaired expression of antiviral genes. Dexamethasone also acted downstream of IFNβ to inhibit expression of a subset of interferon-regulated genes. We tested a number of hypotheses based on previous publications, but found that no single mechanism could account for more than a small fraction of the broad suppressive impact of dexamethasone on macrophage type I interferon signaling, underlining the complexity of this pathway. Preliminary experiments indicated that dexamethasone exerted similar inhibitory effects on primary human monocyte-derived or alveolar macrophages.

## INTRODUCTION

Since the discovery in the mid 20^th^ Century of the potent anti-inflammatory properties of endogenous glucocorticoid (GC) hormones, synthetic GCs have been very widely used in the treatment of both chronic and acute inflammatory pathologies (1). This is despite the fact that they may cause a variety of adverse effects, some of which can be life-threatening in extreme cases (2). Such adverse effects were recognized very early: indeed several were mentioned by Philip Hench in his Nobel Prize lecture of 1950, just two years after his ground-breaking clinical trial of the GC cortisone. Although it was not mentioned by Hench, immunosuppression is a predictable consequence of the clinical use of GCs (3). Activation of the innate immune system is inseparable from inflammation, hence inhibition of inflammation is often accompanied by impairment of innate immunity.

Effects of natural and synthetic GCs are mediated by the GC receptor (GR), a member of a large family of ligand-activated transcription factors (3). Ligand-bound GR can either activate transcription (most often by binding as a dimer to palindromic GC response elements) or suppress transcription (for example via functional interference with the transcription factor NF-κB). It was initially thought that harmful effects of GCs were mediated by transcriptional activation, and beneficial anti-inflammatory effects by transcriptional repression (transrepression) (4). It is now increasingly recognized that this model is too simplistic. Instead, anti-inflammatory effects of GCs involve both transcriptional activation and repression (5, 6). For example, in several cell types GCs increase the expression of dual specificity phosphatase 1 (DUSP1), which exerts anti-inflammatory effects by inactivating mitogen-activated protein kinases (MAPKs), in particular MAPK p38 (7-11).

Monocyte-derived and tissue resident macrophages play central roles in both chronic inflammatory pathologies such as rheumatoid arthritis (12, 13) and acute inflammatory pathologies such as the viral sepsis that can be unleashed by the zoonotic pathogen SARS-CoV-2 (14, 15). Macrophages are important targets of the beneficial anti-inflammatory effects of endogenous and exogenous GCs (8, 16). To investigate how macrophages respond to GCs we and many others have used the Toll-like receptor 4 (TLR4) agonist lipopolysaccharide (LPS) as a stimulus. This reagent has some unique advantages as an experimental tool. When it engages TLR4 it initiates two distinct signaling responses that drive more or less discrete programmes of gene expression (17). At the cell surface, TLR4 uses the adaptor molecule MyD88 (myeloid differentiation primary response 88) to activate NF-κB and mitogen-activated protein kinase cascades. These cooperate to drive rapid expression of several pro-inflammatory genes. Following internalization to endosomes, TLR4 also signals via the adaptor TICAM1 (TIR domain containing adaptor molecule 1) to activate TBK1 (Tank-binding kinase 1) promoting phosphorylation of the transcription factor IRF3 (interferon response factor 3), which in turn induces expression of the type I interferon IFNβ. Macrophages express the dimeric IFNβ receptor encoded by *Ifnar1* and *Ifnar2* genes, and can therefore respond in autocrine or paracrine fashion to secreted IFNβ. This causes activation of the tyrosine kinases JAK1 (Janus kinase 1) and TYK2 (tyrosine kinase 2), which then phosphorylate and activate the transcription factors STAT1 (signal transducer and activator of transcription 1) and STAT2. STAT1 can function as a homodimer to regulate transcription via palindromic GAS elements. STAT1 and STAT2 can also dimerise with one another and combine with a third protein, IRF9, to form the heterotrimeric transcription factor ISGF3 (interferon-stimulated gene factor 3), which binds to a distinct regulatory sequence known as an ISRE (interferon-stimulated response element) (18). ISGF3 and STAT dimers promote expression of hundreds of genes, many of which are mediators of anti-microbial defence, but some of which are pro-inflammatory cytokines and chemokines. The macrophage response to LPS is partly dependent on this positive feedback loop mediated by IFNβ (19–22). The use of LPS as a stimulus therefore allows researchers to examine a broad spectrum of pro-inflammatory and antimicrobial macrophage functions. Moreover, TLR4 can be activated by several DAMPs (damage-associated molecular patterns) (23), as well as by viral proteins like the SARS-CoV-2 spike protein (24–26). The pathways and molecular mechanisms functioning downstream of TLR4 are therefore relevant to both infectious and sterile immune pathologies.

Here we set out to investigate the impact of the synthetic GC Dexamethasone (Dex) on macrophage transcriptome and proteome responses to LPS. We also wanted to determine how the phosphatase DUSP1 contributed to effects of Dex. By serendipity, this study revealed an extremely broad effect of Dex on IFNβ signaling in primary mouse bone marrow-derived macrophages, including consistent suppression of many anti-viral factors at both mRNA and protein levels. Dex inhibited both the LPS-induced expression of IFNβ and the regulation of gene expression by IFNβ, but neither of these effects could be explained by existing models of GC action. Our preliminary data indicated that at least some of these phenomena also occurred in primary human monocyte-derived or alveolar macrophages.

## RESULTS

### Dexamethasone broadly inhibits IFNβ-dependent gene expression in LPS-treated macrophages

To investigate the impact of GCs on the macrophage response to LPS, primary mouse bone marrow-derived macrophages (BMDMs) were stimulated with LPS in the absence or presence of a moderate dose of Dex (100nM) for four hours and steady state mRNA abundance was assessed by microarray. The data were first filtered for robust gene induction by LPS (greater than five-fold increase of expression, p < 0.05), leading to the identification of 599 LPS-induced protein-coding transcripts. The responses of these transcripts to the addition of Dex were then assessed. As illustrated by volcano plot (Fig. 1A), the effect of Dex was strongly dominated by inhibition, 169 transcripts being down-regulated and only 13 up-regulated (fold change > 2 and p < 0.05). Cooperative regulation of the *Lcn2* (lipocalin 2) gene by LPS and Dex was previously reported (27) and confirmed here (Fig. 1A).

**Figure 1.**
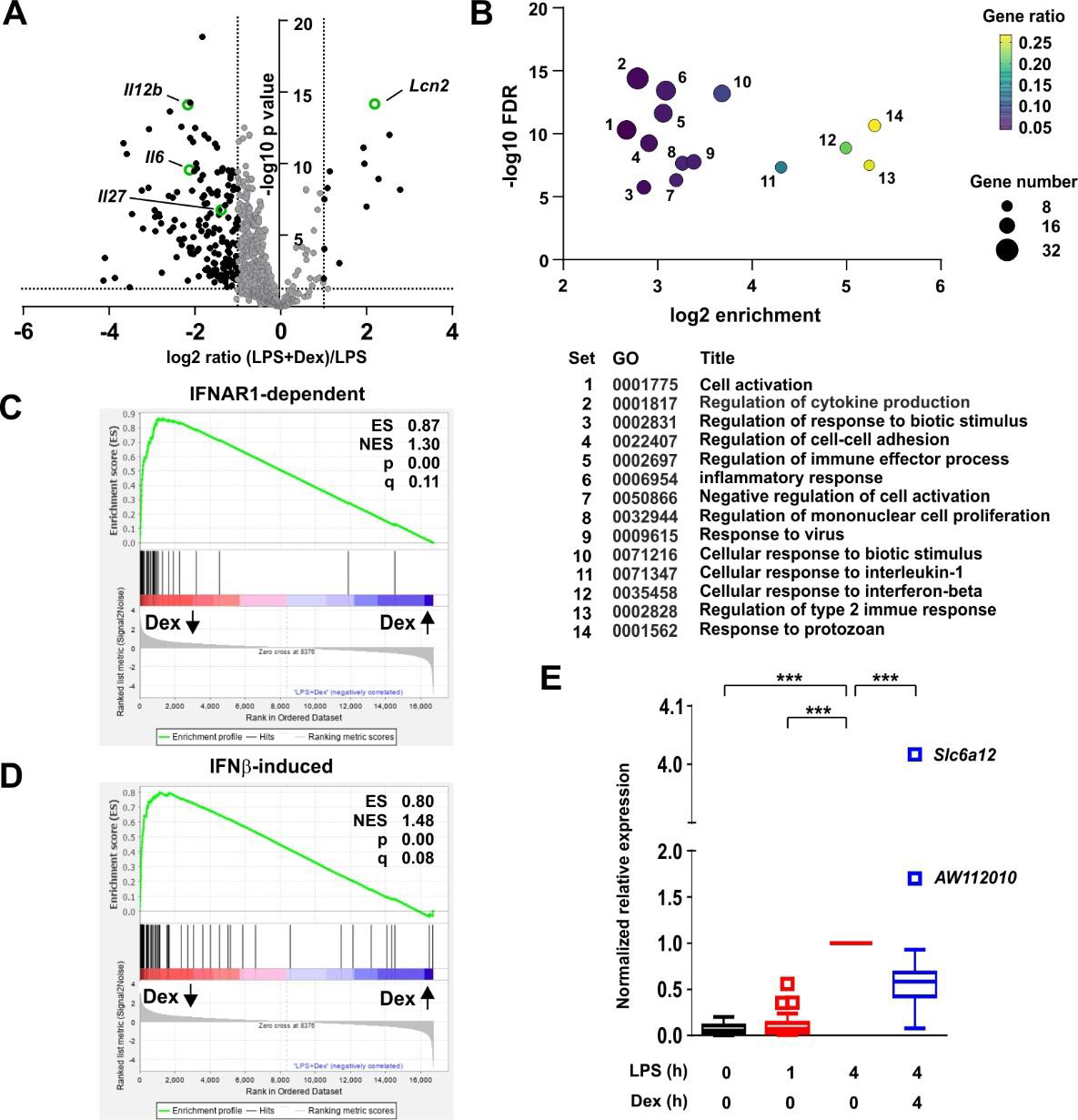
Dexamethasone broadly inhibits IFNβ-dependent gene expression in LPS-treated macrophages. **A)** Mouse BMDMs (three independent isolates per condition) were treated with vehicle, 10 ng/ml LPS for 1 h or 4 h, or 10 ng/ml LPS + 100 nM Dex for 4 h. mRNA was harvested, quantified by microarray and analysed using Partek. Transcripts robustly induced by LPS at 4 h (fold increase > 5, pAdj < 0.05) were identified and their responses to addition of Dex illustrated by volcano plot. Black dots represent LPS-induced transcripts whose expression was significantly altered by addition of Dex (fold change > 2, pAdj < 0.05). The transcripts highlighted in green are discussed in the text. **B)** LPS-induced and Dex-suppressed transcripts (black dots in left quadrant of **A**) were subjected to GO analysis using default parameters in Panther. Significantly enriched GO terms were filtered manually and using REVIGO to eliminate redundant terms. The top 14 remaining terms were plotted according to enrichment, FDR, gene number and gene ratio. **C)** Gene set enrichment analysis was performed using the comparison of LPS vs. LPS + Dex as a quantitative data set, and the top 70 IFNAR1-dependent transcripts from (28) as a categorical gene list. Dex-inhibited transcripts are clustered at the left, and Dex-enhanced transcripts at the right. ES, enrichment score; NES, normalised enrichment score; q, false discovery rate. **D)** Gene set enrichment analysis was performed as in C, except using the 93 most robustly IFNβ-inducible genes from (29). **E)** For 96 genes that were robustly induced by LPS in our data set and interferon-regulated according to (28) or (29) gene expression was normalized to the 4 h value, and mean expression was plotted in Tukey box-and-whisker format. Outliers exceed the median by more than 1.5x interquartile range. ***, p < 0.005.

Gene Ontology (GO) analysis of GC down-regulated genes revealed highly significant enrichment of terms related to inflammation, cytokine production and responses to biotic stimuli (Fig. 1B, Supplemental Table 1). Enrichment of such terms was predictable and uninformative, since the original set of 599 transcripts was selected on the basis of strong induction by LPS. Less predictable were the gene sets “Response to virus” (GO:0009615; set 9 in Fig. 1B) and “Cellular response to interferon beta” (GO:0035458; set 12 in Fig. 1B), which displayed 10.4-fold and 31.8-fold enrichment, respectively. Although only four genes appeared in both lists (Supplemental Table 1), both GO terms allude to antiviral programs of gene expression. Closer inspection revealed many interferon-stimulated genes (ISGs) amongst the LPS-induced genes that were significantly down-regulated by Dex. To explore this in more detail we made use of two published data sets related to interferon-mediated gene regulation in primary murine macrophages (generated in a similar manner). First, a systems-based analysis of the macrophage transcriptome identified a large number of genes whose delayed induction by lipidA (a highly TLR4-specific LPS moiety) was dependent on autocrine signaling via the interferon receptor IFNAR1 (28). The top 70 IFNAR1-dependent genes from that study were used as a gene set for GSEA (gene set enrichment analysis) (Fig. 1C). This confirmed extremely strong enrichment of IFNAR1-dependent genes amongst those negatively regulated by Dex in our own data set, with an enrichment score of 0.866 and p value 0.0. From a study of macrophage regulatory networks (29) we then identified genes robustly induced by treatment of BMDMs with IFNβ alone (fold change > 100 and p < 0.05), and used this list of 93 genes in GSEA (Fig. 1D). Once again there was very strong enrichment of IFNβ-inducible genes amongst those suppressed by Dex in our microarray set, with enrichment score 0.800 and p = 0.0. Removal of genes common to the two lists reduced the enrichment score only marginally (ES = 0.775) and did not change the nominal p value of zero.

We generated a list of 96 genes that were robustly induced by LPS in our own data set and interferon-regulated according to one or both of the above-mentioned data sets (28, 29). Mean responses of these genes to LPS and/or Dex were then plotted (Fig. 1E). The great majority were expressed in a delayed fashion, with little or no increase of expression 1 h after addition of LPS (Fig. 1E). At the 4 h time point, expression of 86 out of 92 genes was significantly inhibited by Dex. Importantly, the inhibitory effect of Dex was not quite universal. The IFNβ-responsive gene *Acod1* (aconitate decarboxylase 1), previously known as *Irg1* (immune-responsive gene 1), mediates production of the anti-inflammatory metabolite itaconic acid (30, 31). The *Acod1* transcript was induced more than 1000-fold by LPS, and this response was spared from inhibition by Dex. Two IFNβ-responsive genes were significantly cooperatively regulated by LPS and Dex (the outliers *AW112010* and *Slc6a12* in Fig. 1F: p = 1.73×10^−4^ and p = 1.00 x 10^−7^, respectively).

Many of the Dex-sensitive genes identified in the above approaches were antiviral mediators. We therefore set out to investigate the extent of the impact of Dex on the macrophage antiviral program by identifying genes that were induced by LPS (log_2_ fold change > 1, p < 0.05); expressed in a delayed manner (at least two-fold higher at 4 h than at 1 h); inhibited by Dex (p < 0.05); labelled as type I interferon-inducible in mouse according to the Interferome database (32); and had plausible roles in the regulation of viral life cycles (at least two supporting publications found in Pubmed using the systematic gene name and “antiviral” as search terms). This generated a list of 107 genes (Supplemental Table 2), which is likely to be under-inclusive because of the use of a single, relatively early time point for analysis of gene expression; shortcomings of the text-mining approach; or shortcomings in functional annotation of genes. For example the poorly annotated gene *AA467197*, mouse ortholog of the human *C15orf48* gene, has only recently been identified as an interferon regulated gene and a putative antiviral factor (33, 34). It is also important to note that viruses may co-opt cellular antiviral machinery to gain a competitive advantage, therefore not all of the genes in the final list are unequivocally “antiviral” under all circumstances. However, many of these are known to contribute to cell-intrinsic defence against SARS-CoV-2 and other viral pathogens (35, 36). These genes belong to several classes including: GTPases or GTP-binding proteins, which contribute to antimicrobial defence via a number of mechanisms (37) (*Gbp2*, *Gbp4*, *Gbp5*, *Gbp9*, *Gnb4*, *Iigp1*, *Irgm2*, *Mx1*, *Mx2*, *Tgtp1*, *Tgtp2*); PRRs involved in the recognition of intracellular pathogens and the initiation of antiviral responses (*Aim2*, *Ddx58* [RIG-I], *Dhx58*, *Eif2ak2* [PKR], *Ifi203*, *Ifi204*, *Ifih1* [MNDA-5], *Mndal*, *Nlrc5*, *Pyhin*); several members of the poly-ADP ribosyltransferase (PARP) and Schlafen (SLFN) families that contribute to antiviral defences (38, 39); and diverse restriction factors that act at distinct points of infectious cycles to prevent entry, replication or exit of viruses (*Bst2* [tetherin], *Ch25h*, *Cmpk2*, *Herc6*, *Ifi35*, *Ifi44*, *Ifi47*, *Ifit1*, *Ifit2*, *Ifit3*, *Isg15*, *Isg20*, *Oasl1*, *Rsad2* [viperin], *Trim21*, *Usp18*) (35). Patterns of expression of the top 50 most Dex sensitive genes are illustrated in heatmap form in Fig. 2A, whilst the full list is included in Supplemental Table 2.

**Figure 2.**
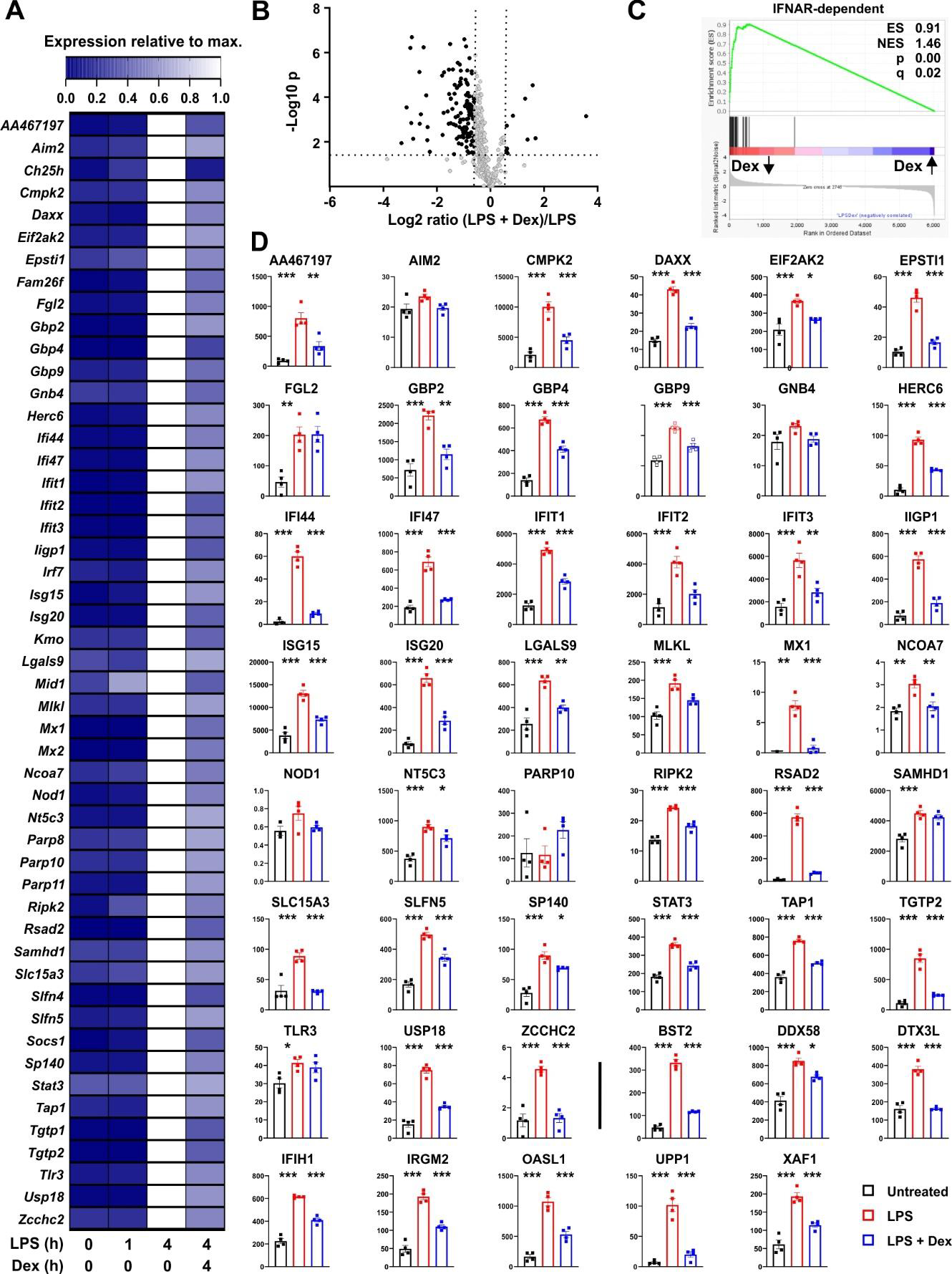
Dexamethasone inhibits expression of multiple antiviral mediators at both mRNA and protein levels. **A)** To identify putative interferon-regulated antiviral mediators that are suppressed by Dex we filtered our data set for genes that were 1) well-annotated; 2) robustly induced by LPS (Log_2_ FC > 1, padj < 0.05); 3) expressed in delayed manner (mean expression at least 2-fold higher at 4h than at 1 h); 4) significantly down-regulated by Dex (padj < 0.05); 5) annotated as type I interferon regulated genes according to the Interferome database (32); 6) with plausible evidence of a role in the regulation of viral life cycles (at least two supporting publications found by PubMed search with appropriate gene symbol and “antiviral”). For these 107 genes (See Supplemental Table 2), mean expression was normalized to expression after 4 h stimulation with LPS. The 50 most Dex-sensitive genes are illustrated in heat map format, ordered alphabetically. **B)** Mouse BMDMs (four independent isolates per condition) were treated with vehicle, 10 ng/ml LPS or 10 ng/ml LPS + 100 nM Dex for 24 h. Protein lysates were generated and subjected to data independent acquisition-based quantitative proteomic analysis. The 410 proteins whose levels were increased by LPS treatment (> 1.5 fold change, padj < 0.05) are illustrated in volcano plot according to their response to addition of Dex. Black dots indicate proteins that whose levels were significantly affected by Dex (> 1.5 fold increase or decrease, padj <0.05). **C)** Gene set enrichment analysis was performed essentially as in 1C. Protein expression data were clustered according to response to Dex, with Dex-suppressed proteins on the left; protein accession numbers were substituted by corresponding gene symbols; and IFNAR-dependent genes from (28) formed the categorical list. **D)** Where data were available from the proteomic study, levels of proteins corresponding to the genes listed in A were plotted. Numerical values are protein concentrations in nM. The last eight plots (after the vertical bar) show proteins of interest, whose corresponding transcripts were not in A. *, p < 0.05; **, p < 0.01; ***, p < 0.005. Absence of symbol indicates lack of statistical significance.

An unbiased proteomic screen was then used to assess protein levels in BMDMs treated with vehicle, LPS or LPS + Dex for 24 hours. Depth of proteome coverage was good, with more than 6,000 proteins detected, and normalised protein concentrations were similar across conditions, indicating an absence of systemic bias (not shown). We identified proteins whose abundance was increased by LPS (> 1.5-fold increase, Padj < 0.05). The lower threshold for fold change was chosen because LPS-induced changes of abundance were generally lower at the protein than at the mRNA level. This generated a list of 410 LPS-regulated proteins, whose responses to Dex were then examined. According to volcano plot, the effect of Dex was again dominated by suppression (Fig. 2B). 125 LPS-induced proteins were significantly negatively regulated by Dex, and 96 of these were identified as type I interferon-regulated genes in the Interferome database (32). Amongst these Dex sensitive targets, gene ontology analysis revealed very strong enrichment of terms related to type I interferons (response to interferon-beta, GO:0035456, fold enrichment 41.2, p = 3.2 x 10^−19^; cellular response to interferon-beta, GO:0035458, fold enrichment 40.9, p = 9.3 x 10^−17^). Gene set enrichment analysis also revealed exceptional enrichment of IFNAR-dependent genes that were negatively regulated by Dex at the protein level (Fig. 2C). Where data were available in the proteomic data set, we examined the expression of individual protein products of the top 50 Dex-sensitive genes (Fig. 2D). With a few exceptions, patterns of expression were similar at mRNA and protein levels, with significant increase in response to LPS and significant decrease in response to addition of Dex. The extent of agreement is striking given that mRNAs were sampled at 4 and proteins at 24 hours after stimulation. The selection of the 24 h time point for assessment of protein expression may have led us to miss or underestimate effects of both LPS and Dex in this experiment. Nevertheless, the emerging picture is one of strong and sustained GC-mediated suppression of antiviral programs. Note that some mediators of antiviral responses (eg. STING1 and MAVS) were constitutively expressed at both mRNA and protein levels and neither up-regulated by LPS nor down-regulated by Dex (data not shown).

Several transcripts encoding secreted or cell-surface immunomodulatory factors were also expressed in delayed fashion, negatively regulated by Dex, and identified as ISGs according to the Interferome database (32) (Fig. 3A). *Il12b*, *Il6* and *Il27* mRNAs were highlighted in Fig. 1A. Levels of the corresponding proteins were measured in supernatants of BMDMs treated under the same conditions (Fig. 3B). All of these cytokines displayed no increase of expression 1 h after an LPS stimulus, strong increase at 4h, and strong inhibition by Dex. Some of the immunomodulatory proteins were measurable in the unbiased proteomic screen, up-regulated by LPS and suppressed by Dex (Fig. 3C). Here we also confirmed the strong induction of LCN2 protein by the combination of LPS and Dex (Fig. 3C).

**Figure 3.**
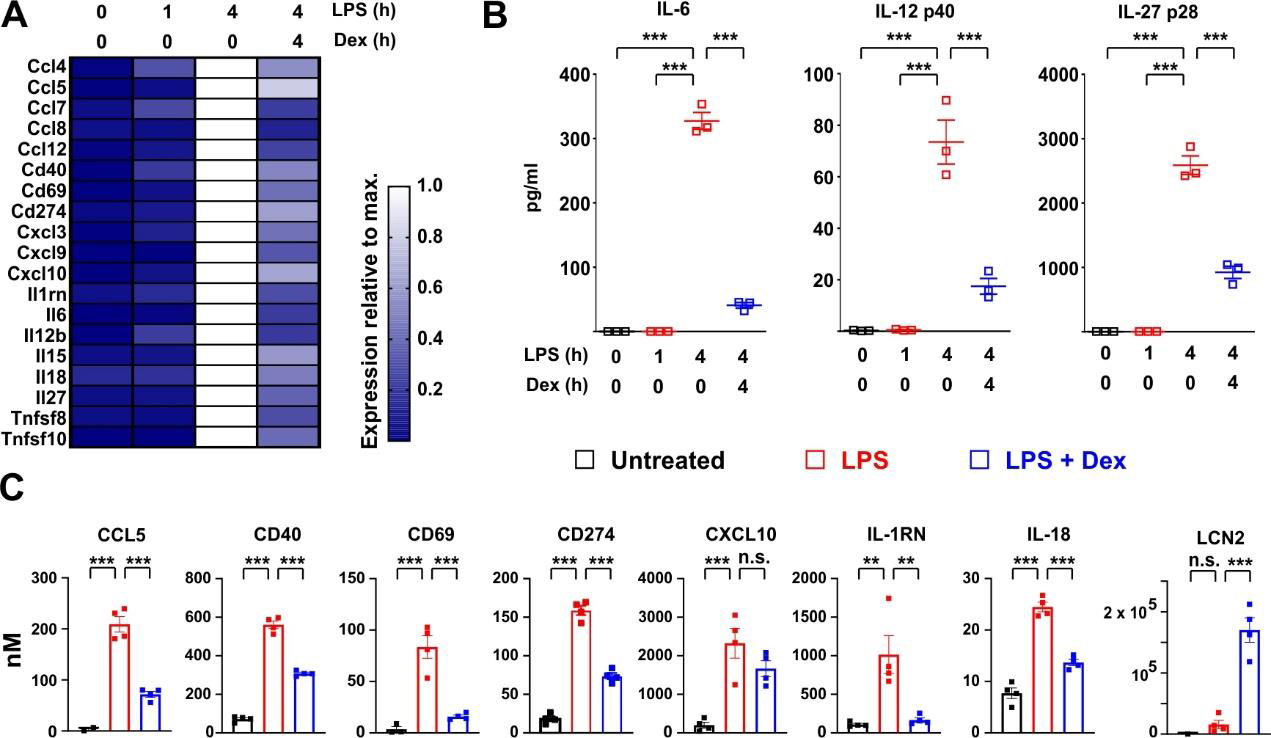
Dexamethasone inhibits expression of multiple cell surface and secreted immuno-modulators at both mRNA and protein levels. **A)** Based on the same microarray data as analysed in Fig.1, expression of selected interferon-regulated immuno-modulatory proteins is illustrated in heat map form as in Fig. 2A. **B)** In a separate experiment mouse BMDMs (three independent isolates) were treated as indicated, and secreted IL-6, IL-12 p40 and IL-27 p28 were measured in supernatants. ***, p < 0.005. **C)** Where data were available from the proteomic study, the expression of immunomodulatory proteins was plotted as in Fig. 2D. **, p < 0.01; ***, p < 0.005; n.s., not statistically significant.

### Dexamethasone strongly inhibits expression of IFNβ by LPS-activated BMDMs

ISGF3 plays a central role in the regulation of ISGs (18). Ligand-activated GR was reported to inhibit expression of ISGs by competing with ISGF3 for an essential transcriptional cofactor (40). To investigate the site of action of Dex we therefore used a mouse macrophage cell line stably transfected with an ISGF3-dependent reporter construct. This construct was strongly activated by either LPS or recombinant IFNβ, and in each case the response was ablated by the broad spectrum JAK inhibitor Ruxolitinib, consistent with dependence on the phosphorylation of STAT1 and STAT2 (Fig. 4A). LPS-induced activation of the reporter was dose-dependently inhibited by Dex, whereas the IFNβ-induced response was insensitive to even the highest dose of Dex used (1 μM), suggesting that Dex does not directly interfere with ISGF3 function.

**Figure 4.**
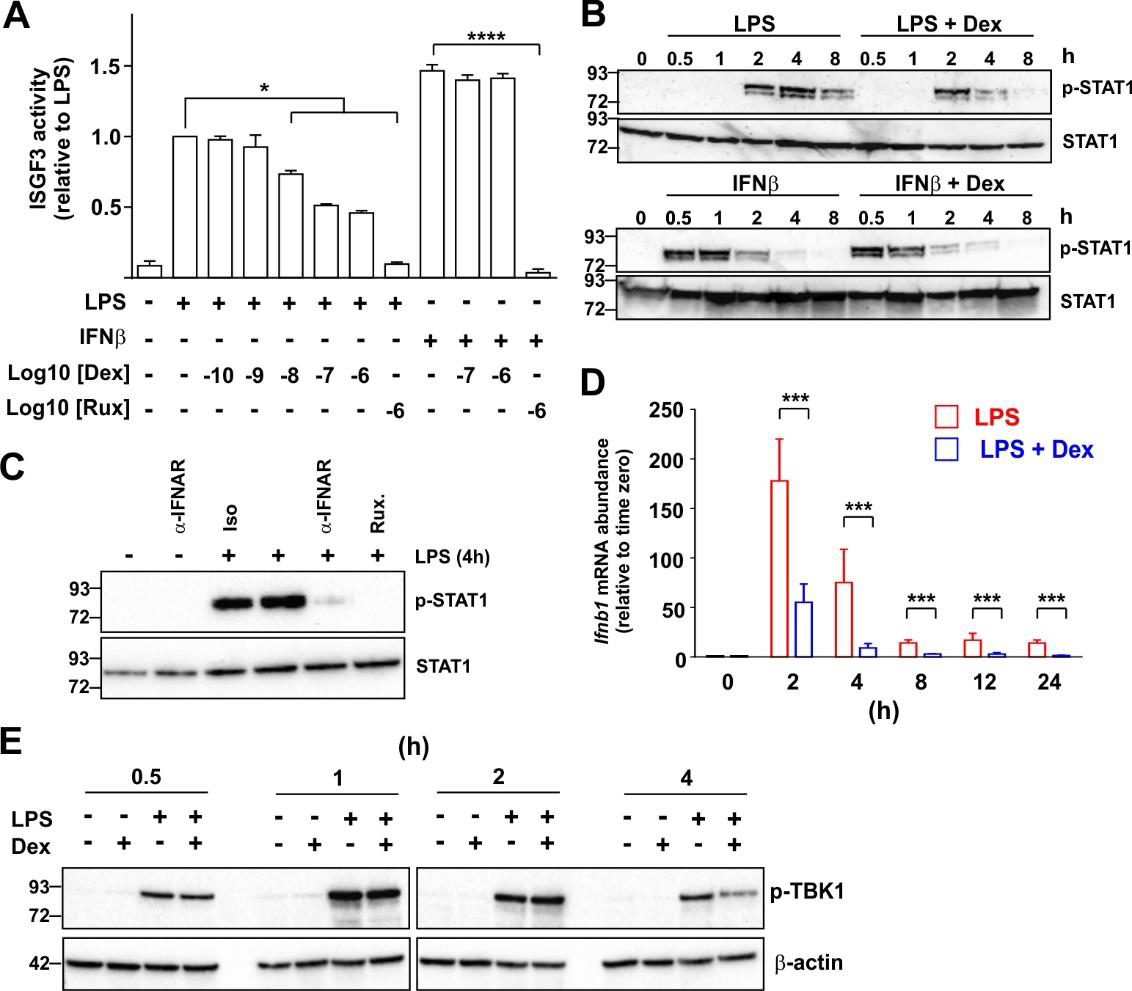
Dexamethasone inhibits expression of IFNβ in LPS-activated mouse macrophages via an unknown mechanism. **A)** A RAW264.7 cell line stably transfected with an ISGF3-dependent SEAP (secreted alkaline phosphatase) reporter was treated with combinations of LPS (10 ng/ml), IFNβ (10 ng/ml), Ruxolitinib (Rux) and Dex as indicated for 24 h and reporter activity was assayed. The graph shows mean results from 3-6 independent experiments ± SEM, with normalization against reporter activity in the presence of LPS alone. *, p < 0.05 (Wilcoxon signed rank test); ***, p < 0.005 (ANOVA). **B)** Mouse BMDMs were treated with combinations of LPS (10 ng/ml), IFNβ (10 ng/ml) and Dex (100 nM) as indicated. Cell lysates were generated and western blotted using antibodies against phosphorylated (Tyr 701) or total STAT1. Representative of three independent experiments. **C)** Mouse BMDMs were stimulated with LPS for 4 h in the absence or presence of an IFNAR neutralizing antibody or isotype control (both 10 μg/ml). Cell lysates were prepared and western blotted as in B. Representative of three independent experiments. **D)** Mouse BMDMs were treated with LPS (10 ng/ml) or LPS + Dex (100nM) for the indicated times, and *Ifnb1* mRNA was measured by qPCR. Mean of three independent experiments ± SEM. ***, p < 0.005. **E)** Mouse BMDMs were treated with combinations of LPS (10 ng/ml) and Dex (100 nM) for 0.5 – 4 h, cell lysates were prepared and blotted for phosphorylated (activated) TBK1 or β-actin. Representative of two independent experiments.

IFNβ caused a rapid increase in the phosphorylation of STAT1, which was evident by 30 minutes and insensitive to Dex (Fig. 4B). In contrast, LPS caused delayed STAT1 phosphorylation, which was not evident until 2 h, and was impaired by Dex, particularly at later time points. The delayed activation of STAT1 in response to LPS was dependent on IFNAR1-mediated signaling, and could be prevented by either an IFNAR1-neutralising antibody or treatment of cells with Ruxolitinib (Fig. 4C). These observations suggest that Dex acts upstream of IFNβ biosynthesis to disrupt the IFNβ-mediated autocrine loop that controls macrophage functions. In fact, the *Ifnb1* gene appeared in several of the GO terms discussed above (Supplemental Table 1), and its expression was strongly inhibited by Dex in the microarray experiment (see below). Quantitative PCR confirmed rapid, strong and sustained induction of the *Ifnb1* gene following LPS treatment of BMDMs (Fig. 4D). Throughout a 24 h time course, this response to LPS was inhibited at least 70% by 100 nM Dex.

It was previously reported that pre-treatment of the myeloid cell line U937 with Dex impaired the LPS-induced phosphorylation and activation of TBK1 (41). We therefore considered the hypothesis that inhibition of TBK1 function explains the inhibitory effect of Dex on IFNβ-mediated feedback in primary macrophages. However, simultaneous treatment of primary BMDMs with LPS and Dex caused no impairment of early LPS-induced TBK1 phosphorylation under conditions that resulted in strong inhibition of both *Ifnb1* and ISGs (Fig. 4E). Dex weakly but consistently reduced TBK1 phosphorylation at the four hour time point. However, this late effect was unlikely to contribute to changes in expression of *Ifnb1*, which were already strongly declining by four hours.

Glucocorticoids selectively inhibit the expression of certain pro-inflammatory genes by increasing and prolonging the expression of DUSP1 and inhibiting MAPK p38 (7, 9, 10, 42). DUSP1 is a negative regulator of *Ifnb1* gene expression (43), and both serum IFNβ and hepatic IFNβ-dependent genes were over-expressed in *Dusp1-/-* mice following infection with *E. coli* (44). These observations suggested the hypothesis that Dex impairs expression of IFNβ and IFNβ-dependent genes in LPS-activated BMDMs via the induction of DUSP1. This hypothesis was tested using macrophages isolated from *Dusp1-/-* mice. IFNβ biosynthesis was elevated in *Dusp1-/-* BMDMs as previously reported (43), but similarly inhibited by Dex in *Dusp1+/+* and *Dusp1-/-* BMDMs (Fig. 5A: note different scales of y axes).

**Figure 5.**
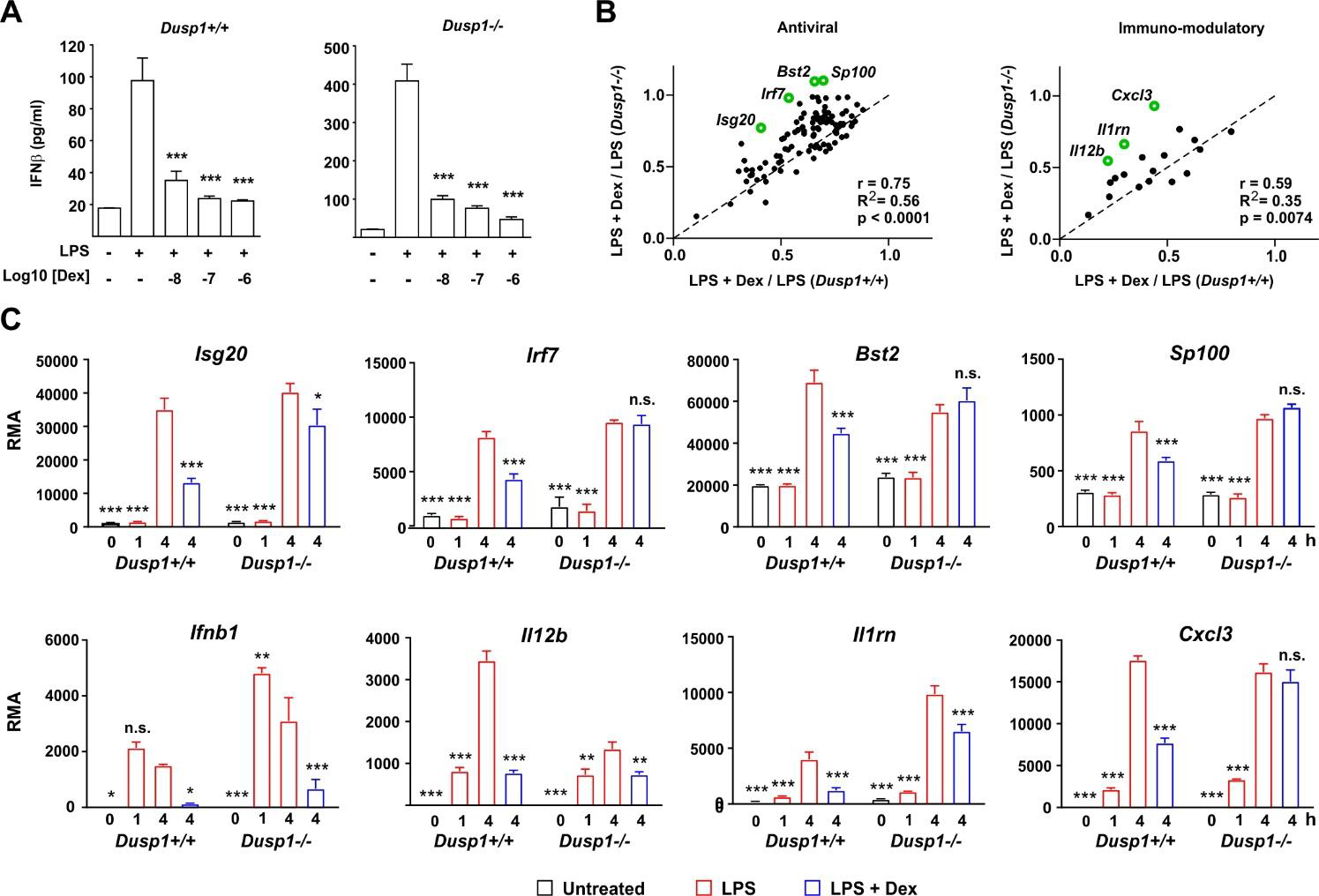
DUSP1 contributes to Dex-mediated suppression of ISGs. **A)** *Dusp1+/+* or *Dusp1-/-* BMDMs (three independent isolates of each) were treated with LPS (10 ng/ml) for 4h with or without addition of Dex as indicated. IFNβ protein was measured by Luminex assay. The graph shows mean ± SEM, n = 3. ***, p < 0.005. **B)** For each interferon-regulated gene of “antiviral” or “immmuno-modulatory” subsets (see Figs. 2 and 3), Dex sensitivity was calculated as the ratio of expression in the presence of LPS + Dex vs. expression in the presence of LPS alone. In the plot of Dex sensitivity in *Dusp1+/+* and *Dusp1-/-* BMDMs, the dotted diagonal represents the null hypothesis, that effects of Dex are independent of *Dusp1* genotype. Selected outlier genes that do not fit the null hypothesis are highlighted. **C)** Microarray-derived expression data for these outlier genes are illustrated. The pattern of expression of *Ifnb1* itself is also shown. *, p < 0.05; **, p < 0.01; ***, p < 0.005; all in comparison to 4 h LPS treatment.

We then systematically examined the impact of DUSP1 depletion on the Dex sensitivity of IFNβ-regulated antiviral genes (Fig. 5B, left) and immunomodulatory genes (Fig. 5B, right). For the gene sets discussed above, Dex sensitivity was calculated as average ratio of expression under conditions (LPS + Dex / LPS). In plots of Dex sensitivity in *Dusp1+/+* (x axis) and *Dusp1-/-* (y axis) backgrounds the dotted diagonal line represents the null hypothesis that disruption of the *Dusp1* gene has no impact on responsiveness to Dex. For both gene sets, sensitivity to Dex was closely correlated in *Dusp1+/+* and *Dusp1-/-* BMDMs, supporting the null hypothesis and suggesting a minimal impact of DUSP1 depletion. However, there were several outliers whose response to Dex was altered by *Dusp1* gene disruption. Some of these are highlighted in Fig. 5B, and primary data from the microarray experiment are illustrated in Fig. 5C. In the cases of *Irf7*, *Bst2*, *Sp100* and *Cxcl3*, significant inhibitory effects of Dex were completely lost in the absence of DUSP1. We conclude that overall DUSP1 plays a minor and highly gene-specific role in the regulation of IFNβ signaling by Dex. Fig 5C also illustrates the behavior of the *Ifnb1* gene itself in the microarray experiment, confirming elevated expression in *Dusp1-/-* BMDMs and strong suppression by Dex in BMDMs of both genotypes.

### Dexamethasone selectively inhibits IFNβ-induced gene expression in BMDMs

Collectively, these observations led to a clear and testable hypothesis: that Dex impairs the responses of macrophages to LPS by inhibiting the expression of IFNβ and disrupting the IFNβ-mediated autocrine feedback loop. We predicted that secondary genes, which are induced by LPS in an IFNβ-dependent manner, would be insensitive to Dex if activated directly by IFNβ, by-passing the site of GC action. To test this hypothesis we first generated a panel of test genes comprising three intracellular antiviral effectors (*Ifit2*, *Rsad2* and *Iigp1*) and three secreted cytokines or chemokines (*Cxcl9*, *Il6* and *Il27*). IFNAR1 dependence of all of these genes was confirmed by reference to a published data-set (28); by our own experiments using *Ifnar1-/-* BMDMs (data not shown); and by use of an IFNAR1-blocking antibody (Fig. 6A). We then tested the responses of all six genes to challenge with LPS or IFNβ in the absence or presence of Dex (Fig. 6B). All three of the antiviral effector genes responded as predicted by the hypothesis: their expression in response to LPS was significantly impaired by Dex, whereas their expression in response to IFNβ was insensitive to Dex. Surprisingly, all three of the cytokine/chemokine genes were suppressed by Dex whether induced by LPS or IFNβ. Dex may therefore act both upstream and downstream of IFNβ to impair LPS responses of BMDMs. The IFNβ-induced expression of *Ifit2*, *Rsad2*, *Iigp1*, *Il6* and *Cxcl9* was very strongly dependent on ISGF3 (at least 100-fold lower expression in IFNβ-stimulated *Irf9-/-* BMDMs) (45). *Ifit2*, *Rsad2* and *Iigp1*were insensitive to Dex, whilst *Il6* and *Cxcl9* were suppressed by Dex in IFNβ-stimulated BMDMs. This confirms the conclusion from the reporter gene assay (Fig. 4A); that Dex does not regulate IFNβ signaling in BMDMs via broad suppression of ISGF3 function.

**Figure 6.**
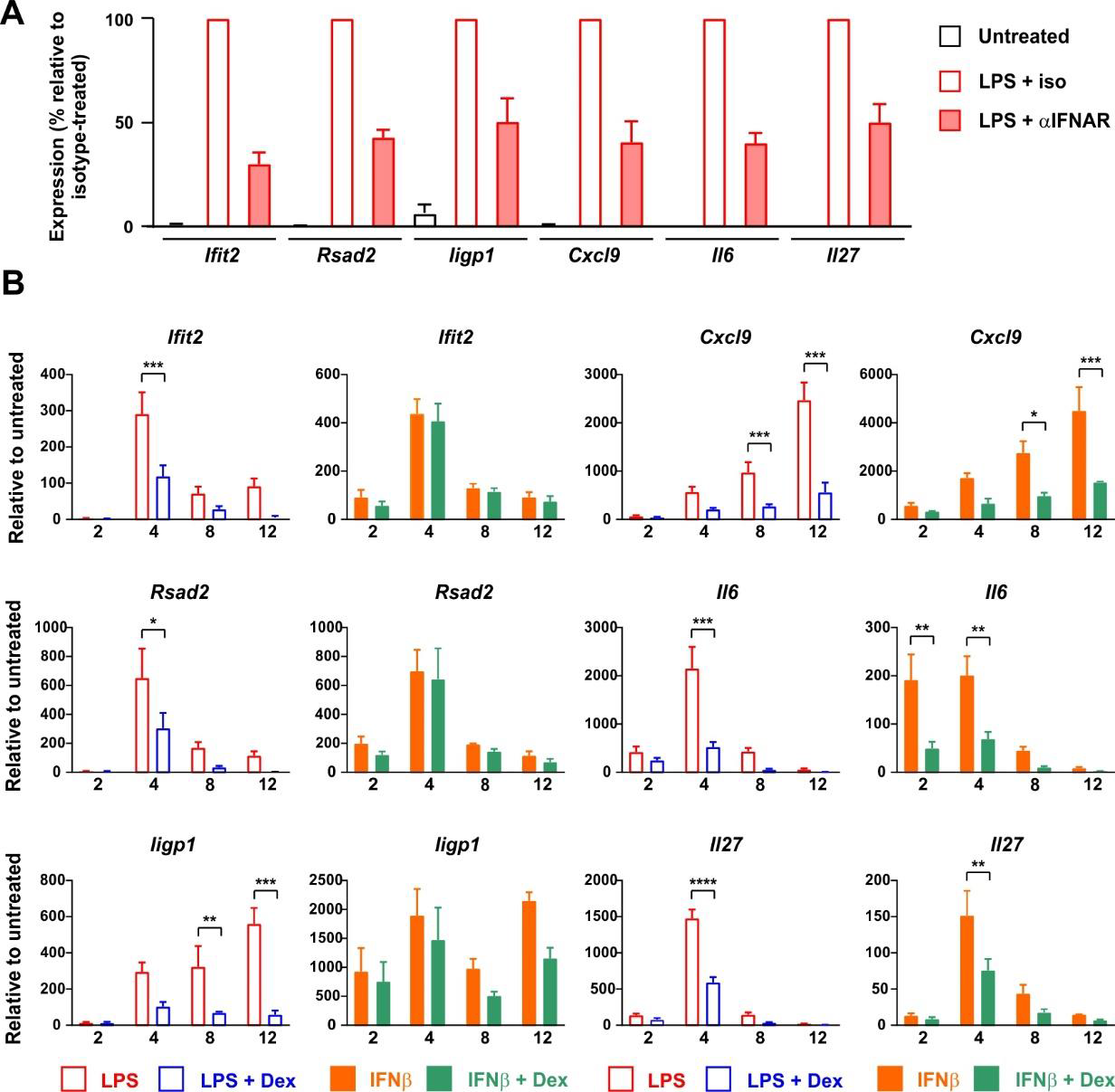
Dex regulates expression of ISGs both upstream and downstream of IFNβ in mouse BMDMs. **A)** BMDMs (4-5 independent replicates) were treated for 4 h with vehicle, 10 ng/ml LPS + 10 μg/ml IFNAR neutralizing antibody, or 10 ng/ml LPS + 10 μg/ml isotype control. RNA was isolated and the indicated transcripts were measured by qPCR, with normalization against LPS + isotype control. **B)** BMDMs (at least three independent replicates) were treated for 0 - 12 h with LPS (10 ng/ml) ± Dex (100 nM) or IFNβ (10 ng/ml) ± Dex (100 nM). Expression of selected genes was determined by qPCR. *, p < 0.05; **, p < 0.01; ***, p < 0.005; absence of symbol indicates lack of statistical significance (p > 0.05).

### Dexamethasone also acts both upstream and downstream of IFNβ in primary human macrophages

To investigate the same phenomena in human cells we isolated monocyte-derived macrophages from peripheral blood of healthy donors and stimulated them with LPS in the absence or presence of 100 nM Dex. Surprisingly there was greater than ten-fold donor-to-donor variation in quantity of IFNβ secreted by LPS-activated MDMs (note logarithmic axis in Fig. 7A). However, IFNβ release was consistently inhibited by 100 nM Dex (mean inhibition ± SEM 52% ± 6%, p = 0.001). Basal expression of some ISGs varied between donors by several hundred-fold (not shown), and magnitudes of response to both LPS and IFNβ were also highly variable (Fig. 7B). This created a practical problem in generating adequate statistical power to investigate effects of Dex, power calculation suggesting that 100 or more individual donors would be needed for some ISGs. Nevertheless a preliminary analysis using seven independent isolates of MDMs confirmed that Dex can inhibit expression of the ISGs *RSAD2* and *CXCL9* whether evoked by either LPS or IFNβ. In human alveolar macrophages the LPS-induced expression of *IFNB1* and *RSAD2* genes was consistently inhibited by Dex (Fig. 7C); however IFNβ protein was below the limit of detection in supernatants of these cells. Scarcity of samples, low yields of mRNA and lack of statistical power prevented us from carrying out an extensive survey of ISG expression. Yet these preliminary data suggest that Dex may also impair type I interferon signaling in alveolar macrophages.

**Figure 7.**
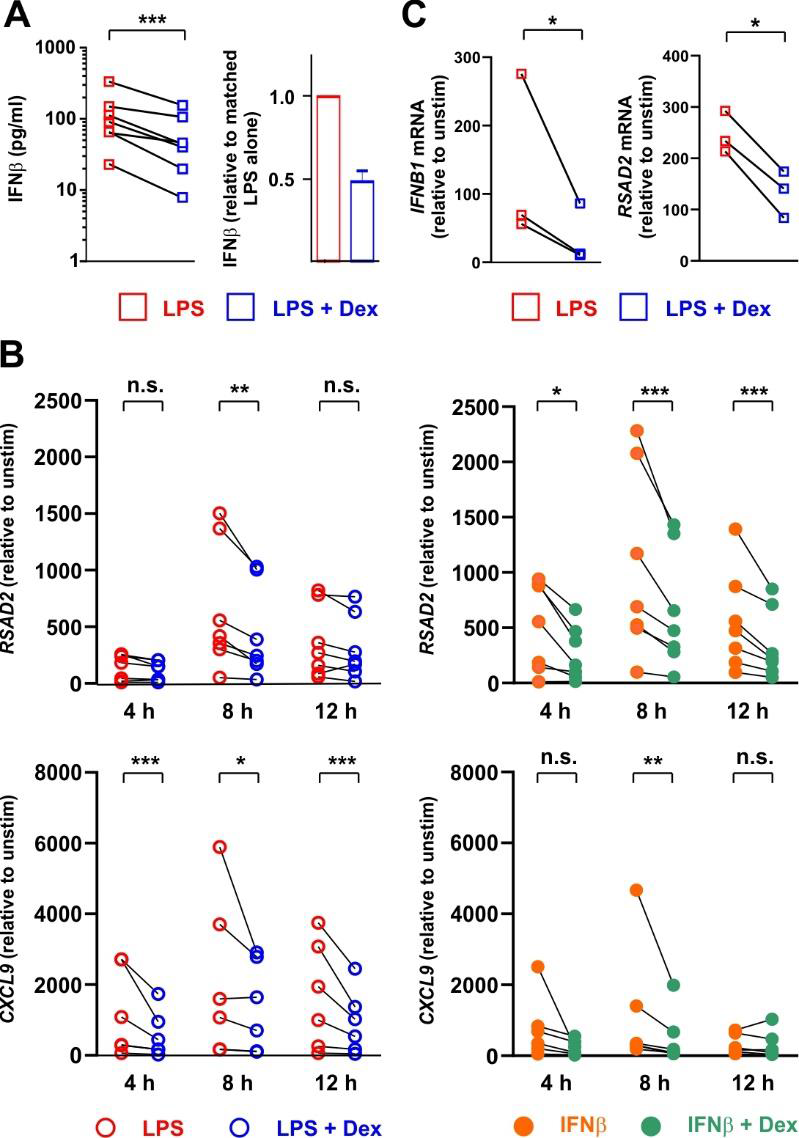
Dex regulates expression of ISGs both upstream and downstream of IFNβ in primary human macrophages. **A)** Primary human monocyte-derived macrophages (n=7) were stimulated with 10 ng/ml LPS ± 100 nM Dex for 4 h, and IFNβ protein in supernatants was measured by Luminex assay. Right hand panel illustrates the same data with normalization to LPS alone. **B)** Primary human monocyte-derived macrophages (n=7) were stimulated with 10 ng/ml LPS ± 100 nM Dex (left) or with 10 ng/ml IFNβ ± 100 nM Dex (right) for 4, 8 or 12 h. *Rsad2* and *Cxcl9* mRNAs were measured by qPCR, with normalization against untreated control for each individual macrophage culture. *, p < 0.05; **, p < 0.01; ***, p < 0.005; n.s., p > 0.05. **C)** Alveolar macrophages were isolated from histologically normal regions of lung resection tissue and treated with 1 μg/ml LPS ± 100 nM Dex for 12 h. *IFNB1* and *RSAD2* mRNAs were measured by qPCR, with normalization against untreated control for each individual sample. *, p < 0.05.

## DISCUSSION

Glucocorticoids have been reported to impair type I interferon signaling in macrophages, but these studies tended to document expression of relatively few ISGs at the mRNA level only (40, 41, 46-48). Here we describe a remarkably broad impact of Dex on the expression of ISGs by LPS-activated BMDMs, occurring at both mRNA and protein levels. Several mechanisms have been suggested to mediate inhibitory effects of GCs on type I interferon signaling, but none of them satisfactorily explain the observations described here. **1)** Dex pretreatment impaired the LPS-induced phosphorylation and activation of TBK1 in a myeloid cell line (41). In our hands Dex did not affect LPS-induced TBK1 phosphorylation under conditions where the expression of *Ifnb1* mRNA and IFNβ protein were strongly inhibited. In any case the impairment of IFNβ production only partially accounts for Dex-mediated impairment of ISG expression in LPS-treated BMDMs, because some ISGs were also sensitive to Dex when directly induced by IFNβ itself. **2)** Downstream of the IFNβ receptor, Dex was reported to inhibit ISG expression by reducing the expression of STAT1 (48), or by up-regulating SOCS1 and inhibiting STAT1 phosphorylation (46). In the presence of either LPS or IFNβ, Dex did not influence STAT1 protein expression over 8 hours. Although Dex inhibited the LPS-induced phosphorylation of STAT1 this was almost certainly a consequence of impaired IFNβ secretion since IFNβ-induced STAT1 phosphorylation was insensitive to Dex. Furthermore, Dex reduced rather than increased the LPS-induced expression of *Socs1* mRNA (LPS alone, 17,621 ± 912; LPS + Dex, 6,278 ± 914). **3)** Another proposed mechanism of action of glucocorticoids is the impairment of ISGF3 function, involving competition between GR and ISGF3 for the essential transcriptional cofactor NCOA2 (nuclear receptor coactivator A2) (40). Several lines of evidence argue against such a mechanism here. First, IFNβ-mediated activation of an ISGF3-dependent reporter construct was insensitive to Dex. In contrast the same reporter was sensitive to Dex when activated by LPS, presumably again due to impairment of IFNβ expression. Amongst our Dex-sensitive ISGs, just over half were reported to have ISGF3 binding sites in their vicinity in a previous study (45) (Supplemental Table 2). However there was poor correlation between ISGF3-dependence in that study and Dex sensitivity in ours (R^2^ = 0.09). The discrepancy is illustrated by the LPS-induced gene *Oas2* (oligoadenylate synthase 2), which was exceptionally dependent on ISGF3 (45) but unaffected by Dex in our study. Although we cannot rule out that Dex inhibits some ISGs via impairment of ISGF3 function, this mechanism cannot explain the broad impact of Dex. **4)** We previously reported that Dex-induced expression of the phosphatase DUSP1 contributes to the suppression of certain LPS-induced genes in macrophages (7, 9, 10, 42). However, the sensitivity of most ISGs to Dex was little affected by disruption of the *Dusp1* gene. In only a few cases was there a clear loss of sensitivity to Dex in *Dusp1-/-* BMDMs. If there is a unifying mechanism by which Dex controls the type I interferon response, it remains to be discovered. Any explanation of this phenomenon must account for its selectivity, and the escape of certain well-known ISGs such as *Acod1* and *Oas2* (49).

Tissue resident macrophages and other myeloid cells play complex roles in infectious diseases such as COVID-19 (14, 15, 50). Myeloid cells are thought to be major sources of pro-inflammatory mediators that cause morbidity and mortality in severe COVID-19. These mediators are not strongly expressed by circulating monocytes, suggesting that the culprits are resident cells at the primary site of infection – the airways. Alveolar macrophages express ACE2 and may become infected by SARS-CoV-2, potentially forming an infective reservoir and secreting pro-inflammatory factors (51–55). The capacity of these cells to mount an effective type I interferon response has been disputed (51, 56). However, more than half of the LPS-induced, interferon-regulated and GC-sensitive antiviral genes discussed here were also upregulated in alveolar macrophages of healthy volunteers following inhalation of LPS (57) (Supplemental Table 2). This suggests that type I interferon responses of human alveolar macrophages at least overlap with those of mouse BMDMs. Circulating monocytes do not express ACE2 and are not thought to be infected by SARS-CoV-2 (53). However they may be activated by Coronavirus spike protein via TLR4 (24–26), or by type I interferons. An early and robust type I interferon signature in circulating monocytes is associated with mild disease following SARS-CoV-2 infection (50). On the contrary, low expression of antiviral genes including *CMPK2*, *EPSTI1*, *HERC6*, *IFI44*, *OASL* and *RSAD2* in peripheral blood was associated with worse clinical outcomes in COVID-19 (58). The responses of myeloid cells to endogenous GCs such as cortisol and therapeutic GCs such as Dexamethasone are highly likely to influence several aspects of disease progression or recovery. Airway stromal cells, considered the primary targets of infection, may also be significantly impacted by GCs. For example in A549 airway epithelial cells stimulated for 18 h with IL-1β, the expression of 84 genes was significantly impaired by addition of Dex (fold decrease >2, p < 0.05) (59). The majority of these genes are interferon-regulated genes according to the Interferome database (32), and many are orthologs of genes that we found to be suppressed by Dex in LPS-treated BMDMs (eg. *Cmpk2*, *Herc6*, *Ifi44*, *Ifih1*, *Ifit1*, *Ifit3*, *Isg15*, *Isg20*, *Oasl1*, *Usp18*). It is likely that Dex also disrupts IFNβ-mediated positive feedback loops in these cells.

In the context of infectious disease such as COVID-19 the consequences of GC-mediated impairment of the type I interferon signaling pathway are likely to be both beneficial and detrimental. On the positive side, the suppression of interferon-regulated cytokines and chemokines (Fig. 2) is likely to contribute to anti-inflammatory effects of GCs. For example, IL-27 is an interferon-regulated member of the IL-12 family of dimeric cytokines (60–62) that has been implicated in pathogenesis of sepsis and chronic inflammatory diseases (63, 64). Serum levels of IL-27 were decreased by glucocorticoid treatment in systemic lupus erythematosus (65), although to our knowledge IL-27 has not previously been identified as a glucocorticoid target in macrophages. Elevated levels of IL-27 also contribute to a signature predictive of poor outcome in COVID-19 (66–68). IL-6 is also thought to have a pathogenic role in severe COVID-19 (69). At the time of writing, the IL-6 receptor-targeting antibody tocilizumab has been trialled but not yet approved for treatment of severe COVID-19 (70). Suppression of IL-6, IL-27 and several other interferon-regulated cytokines and chemokines may contribute to beneficial effects of systemic GCs in severe COVID-19.

On the other side of the equation, type I IFN signaling is essential for host defence against viral pathogens such as SARS-CoV-2 (71, 72). Like other viruses, SARS-CoV-2 deploys several mechanisms to subvert IFN signaling in the host (73). In a similar way, GCs impair the expression of several genes that contribute to host defence against SARS-CoV-2 and other viral pathogens. The genes *Ddx58* and *Ifih1* respectively encode RIG-I and MDA5, which are essential for detection of intracellular SARS-CoV-2 (74) and are targeted by the virus as a means of evading interferon responses (75). Expression of both of these genes is negatively regulated by Dex in BMDMs (Fig. 2). Several of the other genes we found to be robustly inhibited by Dex at both mRNA and protein levels are known to function as restriction factors for SARS-CoV-2 (including *Bst2*, *Ifit1*, *Ifit3*, *Isg15*, *Isg20*, *Rsad2*, *Usp18*) (36). Restriction of nucleotide availability is one mechanism for inhibition of viral propagation (76, 77). The genes *Cmpk2*, *Nt5c3* and *Rsad2*, which function in nucleotide metabolism, were all up-regulated by LPS and inhibited by Dex at both mRNA and protein level (Fig. 2). The same behavior was displayed by *Upp1* (uridine phosphorylase 1): we hypothesise that this gene also encodes an interferon-regulated viral restriction factor, as has been shown for the related gene *UPP2* (36). Finally, Dex inhibited the LPS-induced expression of genes involved in immunoproteasome activation and antigen presentation (*Psmb8*, *Psmb9*, *Psmb10*, *Tap1*, *Tap2* several *H2* alleles; Supplemental Table 2 and data not shown), suggesting that it may impair the engagement of the adaptive immune system to combat viral pathogens. Altogether, the effects of Dex suggest a dismantling of cell intrinsic and extrinsic defences against viral pathogens. An acknowledged limitation of the present work is that we have not yet examined effects of Dex on SARS-CoV-2 infection and propagation. This is a focus of ongoing work.

The delicate balance between harmful and beneficial effects of GCs has been vividly illustrated during the COVID-19 pandemic. Endogenous GC excess is a risk factor for COVID-19 (78). Similarly, prolonged use of synthetic GCs for the treatment of immune-mediated inflammatory diseases both increases the likelihood of contracting COVID-19 and contributes to worse outcomes of the disease (79, 80). The unregulated use of GCs for prophylaxis or treatment of early COVID-19 has been associated with increased incidence of mucormycosis, a disease caused by an opportunistic fungal pathogen (81). Yet no other treatment has proven better than GCs for reducing mortality in patients suffering from severe COVID-19 and requiring respiratory support (82, 83). Contrastingly, in mild or early disease there is no evidence for benefit of GCs (82–84), and some suggestion of harmful effects, for example including delays to viral clearance (85, 86). There are clearly many unanswered questions about when and how best to use GCs against SARS-CoV-2 and other viral pathogens. Indeed it has previously been argued that variable outcomes of GC trials in COVID-19 were related to inconsistent timing of delivery (87). The balance between the harm of suppressing antiviral mechanisms in early disease and the benefit of suppressing the cytokine storm in the later phase is precarious. The tipping point between harm and benefit may also differ between individuals. There is clearly a strong case for application of well-informed patient stratification in COVID-19 (88–90). We argue that the effects of GCs on type I interferon-mediated host defences should be part of such a precision medicine approach. Furthermore, there is urgent need for further research on the mechanistic basis of cross-talk between GCs and type I interferon signaling.

## MATERIALS AND METHODS

### Macrophage isolation and culture

All mice were maintained at the Biomedical Services Unit of the University of Birmingham. Animal care and experimental procedures were performed according to Home Office guidelines and approved by the University of Birmingham Local Ethical Review Committee. Bone marrow was isolated from the femurs of humanely culled 6-12 week old wild type C57BL/6J mice, and BMDMs (bone marrow derived macrophages) obtained by differentiation *in vitro* in RPMI 1640 medium with L-glutamine (Gibco Thermo Fisher Scientific 21875034) supplemented with 10% heat-inactivated fetal bovine serum (Sigma F0392) and 50ng/ml recombinant M-CSF (Peprotech 300-25) for 7 days. BMMs were plated at a density of 1 x 10^6^ml in the appropriate cell culture plate at least 1 d prior to stimulation.

The *Dusp1-/-* strain was a generous gift from Bristol-Myers Squibb, and was back crossed to C57BL/6 for ten generations prior to experiments described here.

Human monocytes from healthy blood donors were isolated from leukapheresis blood cones supplied by the National Blood and Transplant Service (ethical approval ERN_16-0191). Monocytes were enriched by negative selection using StemCell RosetteSep Human Monocyte Enrichment Cocktail (StemCell 15068; 75μl per ml of blood) and Ficoll-Paque (VWR 17144003). Cells were differentiated for 7 days in RPMI 1640 medium with L-glutamine (Gibco Thermo Fisher Scientific 21875034) supplemented with 5% heat-inactivated fetal bovine serum (Biosera FB-1001) and 50ng/ml recombinant M-CSF (Peprotech 300-25). Differentiated macrophages were plated at a density of 1 x 10^6^ml in the appropriate cell culture plate at least 1 d prior to stimulation.

Ethical approval was obtained to recruit adult patients scheduled for surgery to remove lung tissue as part of their clinical treatment plan (predominantly lobectomy for lung cancer) at the Thoracic Surgery Unit at the Queen Elizabeth Hospital Birmingham (REC 17/WM/0272). Lung tissue samples distant from any tumor, without macroscopically evident pathology, and surplus to requirement for histopathology, were washed through with 500 ml of sterile 0.9% saline (Baxter, UK) using a 14 gauge needle (Vasofix ®). The washed through lavage fluid was collected and centrifuged at 4°C and 560g for 10 minutes. Cell pellets were resuspended in 10 ml of RPMI1640 containing 10% FBS, and overlaid onto 10 ml of Lymphoprep^TM^ (StemCell) prior to centrifugation at 800g for 30 min at 4°C. The interphase layer containing mononuclear cells was aspirated into a sterile 50 ml tube containing PBS with 10 % FBS, which was then centrifuged at 300g for 10 minutes at 4°C. AMs were resuspended in RPMI1640 containing 10% FBS, 100U/mL penicillin, 100µg/mL streptomycin and 2mM L-glutamine (Sigma-Aldrich), and plated.

Stimulations were carried out in appropriate culture plates using the following reagents and concentrations unless otherwise stated: LPS (Enzo ALX-581-010-L002; 10ng/ml); recombinant human IFNβ (Peprotech 300-02BC; 10ng/ml); recombinant mouse IFNβ (BioLegend 581306; 10ng/ml); Ruxolitinib (Selleck S1378; 1μM); Dexamethasone (Sigma D8893; 100nM). Where stated, cells were incubated for 2 hours with an IFNAR1 blocking monoclonal antibody (Fisher Scientific MAR1-5A3; 10 μg/ml), or mouse IgG1 kappa isotype control, (Fisher Scientific P3.6.2.8.1; 10 μg/ml) prior to stimulations.

### Stable macrophage cell line ISGF3-dependent reporter assay

The RAW-Blue ISG macrophage cell line stably transfected with an ISGF3-dependent reporter construct was cultured and maintained according to the manufacturer’s instructions (RAW-Blue ISG cells, Invivogen raw-isg). ISGF3 reporter activity was determined from cell culture supernatants using QUANTI-Blue detection reagents according to the manufacturer’s instructions (Invivogen).

### Measurement of mRNA

RNA was isolated from mouse BMMs and human monocyte derived macrophages using Norgen Total RNA Purification Plus kit (Geneflow P4-0016) according to manufacturer’s instructions. cDNA was synthesized using the iScript cDNA Synthesis Kit (Biorad 1708891). mRNA was detected by RT-qPCR using SYBR TB Green Premix Ex Taq (Takara RR820W) and primers supplied by Eurofins Genomics or Sigma Aldrich. UBC (human) or B2M (mouse) were used to normalize mRNA measurements via 2^-ΔΔCt method. Primers are listed in Table 1.

**Table 1.**
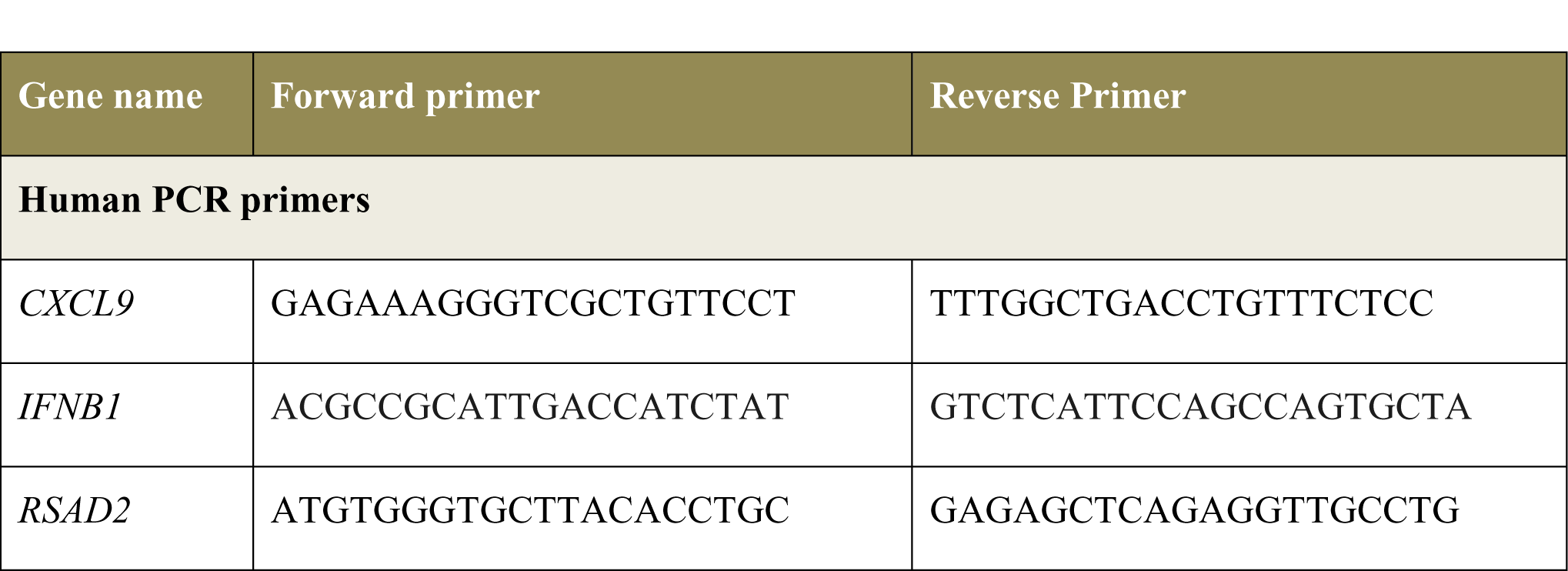
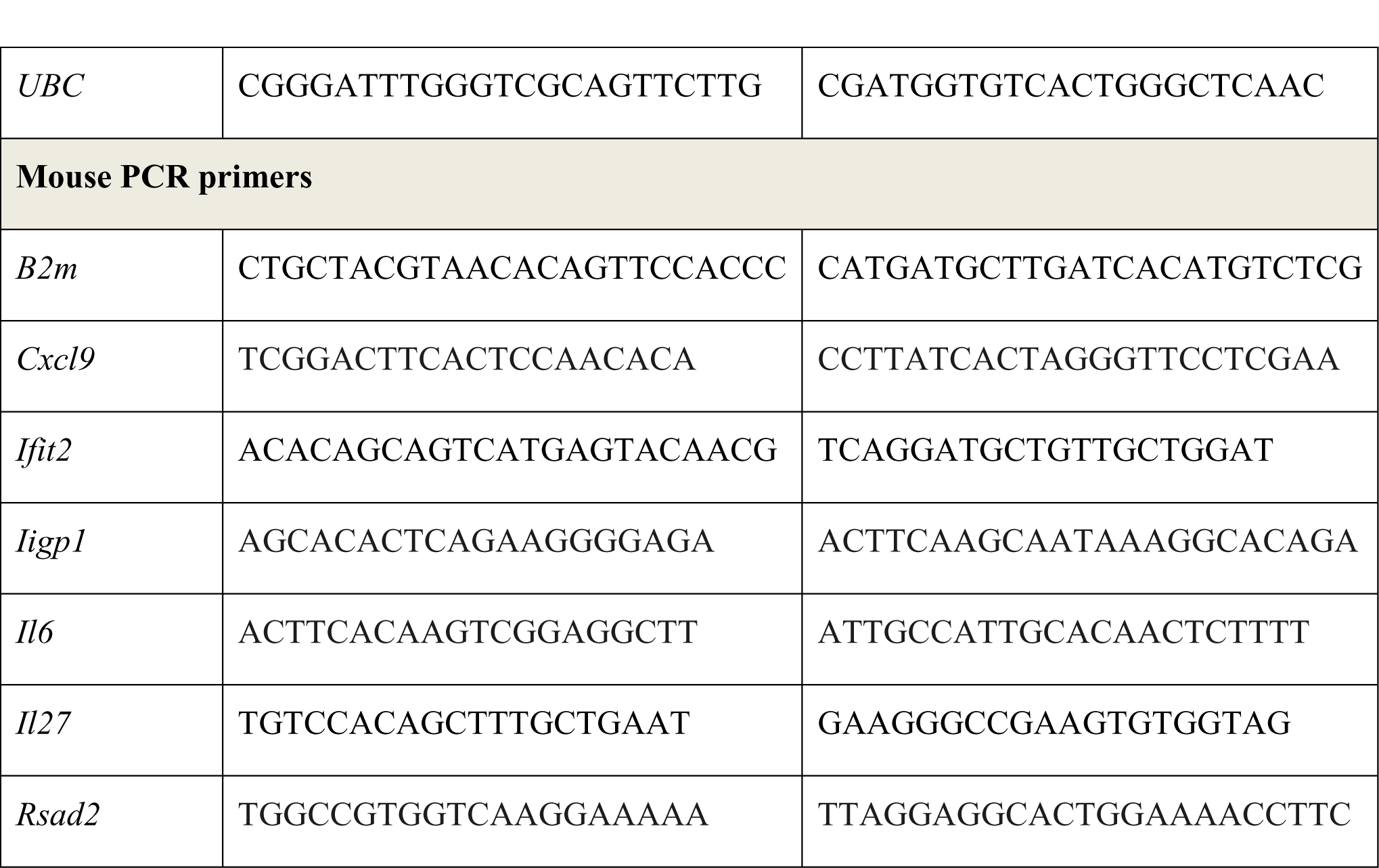
PCR primers used in this study.

### Microarray analysis and bioinfomatics

Microarray analysis was performed using SurePrint G3 Mouse GE 8×60K slides (Agilent) and Partek Genomics Suite version 6.6, build 6.13.0315 (Partek) as previously described (91). For generation of volcano plots, transcripts were first filtered for significant upregulation in response to LPS (> 5-fold increase, adjusted p-value < 0.05) and weakly expressed transcripts were removed by application of an arbitrary filter of 200 RMA. Plots (log_2_ fold difference of expression vs -log_10_ ANOVA p-value) were constructed using Prism (GraphPad Software), with subset cut-offs at p-value < 0.05 and fold difference of expression > 2. Microarray data discussed in this paper were deposited at Gene Expression Omnibus (http://www.ncbi.nlm.nih.gov/geo/) with the accession number GSE68449.

Gene Ontology analysis was performed using the Panther v16 online toolset (92). Redundant GO terms were filtered using REVIGO (93). Gene Set Enrichment Analysis (GSEA) was performed using GSEA_4.1.0 (94).

### Multiplex cytokine analysis

Conditioned medium samples from cultured macrophages were subjected to custom multiplex Luminex assay (Bio-Rad Custom Bio-Plex Assays), and Invitrogen ProcartaPlex Mouse IFNβ simplex, (EXP01A-26044-901), according to manufacturer’s instructions.

### Western blotting

For whole cell lysates, cells were harvested in RIPA buffer and samples passed through a Qiashredder column to disrupt genomic DNA (Qiagen 79656). Protein was quantified by Pierce BCA Assay (Thermo Fisher Scientific 23225). Laemmli or XT sample buffer was added and samples heated to 95°C for 5min. Western blotting was performed using Criterion TGX protein gels (BioRad) and Tris-Glycine SDS buffer (Geneflow B9-0032) or 4–12% Criterion™ XT Bis-Tris Protein Gels (BioRad) and XT MES running buffer (BioRad 1610789). Protein was transferred to BioRad Trans-Blot PVDF membranes (BioRad 1704157) using BioRad Trans-Blot Turbo transfer system. Blots were imaged using Clarity Western ECL Substrate (BioRad 1705061) and BioRad ChemiDoc MP Imaging System. Densitometry for Western blot quantification was performed using ImageJ. Antibodies used in western blotting are listed in Table 2.

**Table 2.** Antibodies used in this study.

| Antibody | Dilution |  | Company |
| --- | --- | --- | --- |
| pSTAT1 | 1:1000 | 58D6 | Cell Signalling |
| Stat1 | 1:2000 | (C-111) sc417 | Santa Cruz |
| B-Actin | 1:5000 | A1978 | Merck |
| pTBK1 / NAK<br>(S172) | 1:1000 | (D52C2) 5483S | Cell Signalling |
| TBK1 / NAK | 1:1000 | (D1B4) 3504S | Cell Signalling |
| IFNAR1<br>Monoclonal | 1:100 (10 ug/mL) | (MAR1-5A3), Cat:<br>16-5945-85 | Fisher Scientific |
| Antibody Functional Grade |  |  |  |
| Mouse IgG1 kappa Isotype Control | 1:50 (10 ug/mL) | (P3.6.2.8.1) Cat: 14-4714-85 | Fisher Scientific |

### Proteomic analysis

Following derivation and stimulation of mouse BMDMs (as described above) cells were washed with PBS and lysed in proteomic lysis buffer (5% SDS, 10mM TCEP, 50mM TEAB). Unbiased proteome analysis was carried out by data-independent acquisition (DIA) mass spectrometry proteomics utilising S-Trap on-column digestion and purification, following the methods detailed by Baker et al. (95). Protein copy number was determined using Perseus software (95, 96) with normalisation for histone protein content. Data were analysed as described in the above methods for microarray analysis and bioinformatics.

### Statistical analysis

GraphPad Prism software (Version 6) was used for statistical analysis. Mann Whitney U test was used for comparison of two groups. For analysis of multiple groups, ANOVA was used with Bonferroni correction for multiple comparisons. The following marks are used throughout: *,p<0.05; **,p<0.01;***,p<0.005; n.s., not statistically significant. N numbers specified in figure legends indicate biological replicates. In human alveolar macrophages, where expression of ISGs was highly variable between donors, ratio paired t test was used to test for consistent effect of Dex.

## AUTHOR CONTRIBUTIONS

AC secured funding for the study. AC, JO, RH and JA contributed to conception and design of the study. JO, OB, SC, TT, KD, SL-R, JW and CM performed experiments and analysed data. RM provided essential clinical samples. All authors contributed to manuscript revisions, read and approved the submitted version.

## DATA AVAILABILTIY

Microarray data discussed in this paper are available at GSE68449. Deposition of proteomic data is in progress at the time of submission.

